# Age differences in retrieval-related reinstatement reflect age-related dedifferentiation at encoding

**DOI:** 10.1101/2020.01.21.912089

**Authors:** Paul F. Hill, Danielle R. King, Michael D. Rugg

## Abstract

Age-related reductions in neural specificity have been linked to cognitive decline. We examined whether age differences in specificity of retrieval-related cortical reinstatement could be explained by analogous differences at encoding, and whether reinstatement was associated with memory performance in an age-dependent or age-independent manner. Young and older adults underwent fMRI as they encoded words paired with images of faces or scenes. During a subsequent scanned memory test participants judged whether test words were studied or unstudied and, for words judged studied, also made a source memory judgment about the associated image category. Using multi-voxel pattern analyses, we identified a robust age-related decline in scene reinstatement. This decline was fully explained by age differences in neural differentiation at encoding. These results suggest that, regardless of age, the specificity with which events are neurally processed at the time of encoding determines the fidelity of cortical reinstatement at retrieval.

## Introduction

The ability to recollect information about a past event declines with advancing age^1–4^. Here, we examined whether age-related differences in recollection are associated with a tendency for older adults to retrieve less detailed or differentiated information about a prior experience than younger individuals. To address this question, we examined age-related differences in retrieval-related *cortical reinstatement* effects. Cortical reinstatement refers to the finding that successful recollection is associated with patterns of cortical activity that partially overlap the patterns elicited when the recollected information was initially experienced (e.g. ^5–7^; for reviews see^8–11^). These findings have led to the widely held view that reinstated cortical activity reflects retrieved content^10^. From this perspective, if there is a tendency to retrieve less detailed or lower fidelity information with increasing age, cortical reinstatement effects should be weaker in older adults relative to younger individuals.

Prior work examining age-related differences in retrieval-related cortical reinstatement effects have reported mixed results^12–19^. However, and of relevance to the present experiment, the analytical approaches employed in the studies that reported null age effects for reinstatement (e.g.^18,19^) characterized reinstatement at the categorical rather than the item level (that is, in terms of patterns of neural activity common to an entire class of encoding events). Thus, these studies leave open the possibility that the fidelity with which finer-grained, item-level information is reinstated differs between young and older adults (see^14,16,17^).

Patterns of neural activity elicited by some stimulus categories become less distinctive in older age. This phenomenon - *age-related neural dedifferentiation* - has been conjectured to play a role in age-related cognitive decline^20–22^. Notably, lower specificity of neural responses to perceptual events has been reported to predict poorer subsequent memory for the events in both young and older adults^23–25^. These findings raise the possibility that the distinctiveness or fidelity with which the perceptual features of an event are neurally represented at the time of encoding (that is, its level of neural differentiation) are a determinant of the fidelity with which the features are reinstated at retrieval.

In apparent contradiction to this possibility, St-Laurent and colleagues^16^ reported that while the viewing of multimodal video clips was associated with only minimal evidence for age-related dedifferentiation, there were pronounced age differences in cortical reinstatement effects as participants mentally ‘replayed’ the videos from memory. These findings were however based on analyses of data that had been pooled across numerous repeated viewings and retrieval attempts, raising the possibility that measures of neural specificity in the young and older adults were differentially influenced by repetition. Indeed, further analysis of the same data-set revealed that neural differentiation of the video clips was greatest for their initial presentation, and declined with subsequent viewings^17^. The authors did not report, however, whether neural differentiation during the initial viewing was associated with the strength or fidelity of neural reinstatement. In a study by Abdulrahman et al.^12^, the authors were also unable to identify any evidence for age-related neural differentiation (operationalized as the accuracy of an MVPA classifier) during the encoding of blocks of words subjected to phonological vs. semantic processing. Robust evidence of age-related reductions in the specificity of reinstatement was nonetheless obtained at retrieval. However, in the retrieval phase of this experiment test items were blocked according to their study context, precluding the analysis of the items according to the accuracy of the associated memory judgment. Thus, it is unclear whether the reported age differences in reinstatement reflected differences in neural correlates of pre- or post-retrieval processing; by definition, reinstatement effects are a reflection only of the latter class of processes.

In the present study, healthy young and older adults encoded words paired with images of faces or scenes as they underwent functional magnetic resonance imaging (fMRI). Face and scene images were selected as stimuli because they have been previously reported to give rise to robust age-related neural dedifferentiation effects (e.g.,^24,26–29^). Studied and unstudied words were subsequently presented in a scanned memory test in which, for words judged as studied, a source memory judgment for the corresponding image category was required. We addressed three principal questions: (1) whether scene and face reinstatement effects associated with successful source memory judgments for the respective stimulus classes differed in their strength or specificity according to age (cf.^12^ vs.^19^), (2) whether age differentially impacts reinstatement of item-vs. category-level information, and (3) whether age differences in retrieval-related reinstatement could be accounted for by analogous differences in neural differentiation at the time of encoding. In further trial-wise analyses, we built on prior findings^30,31^ to ask whether strength of retrieval-related reinstatement covaried within-participants with retrieval-related hippocampal activity, whether either or both of these variables covaried with memory performance, and whether any such relationships differed according to age.

## Results

### Behavioral Results

Means and standard deviations of behavioral measures are presented in Table 1. Vividness ratings and median response times (RTs) from the study phase were sorted according to subsequent memory status into source correct (SC) and incorrect (including source incorrect, ‘Don’t Know’, and item misses) bins and submitted to separate 2 (age) x 2 (memory) x 2 (category) mixed factorial ANOVAs. The analysis of vividness ratings produced a significant main effect of memory (*F*_(1,46)_ = 53.33, *p* = 3.13 × 10^−9^, partial-η^2^ = .54), which was driven by reduced vividness for incorrect memory trials. The main effects of age (*F*_(1,46)_ = 3.12, *p* = .084, partial-η^2^ = .06) and category (*F*_(1,46)_ = 0.70, *p* = .409, partial-η^2^ = .01) were not significant, nor were there any significant interactions involving age or category (all *ps* > .1). The analysis of RTs revealed a significant main effect of category (*F*_(1,46)_ = 5.35, *p* = .025, partial-η^2^ = .10), which reflected faster response times for face trials relative to scenes. The remaining main effects and interactions were not significant (all *ps* > .1).

**Table 1.**
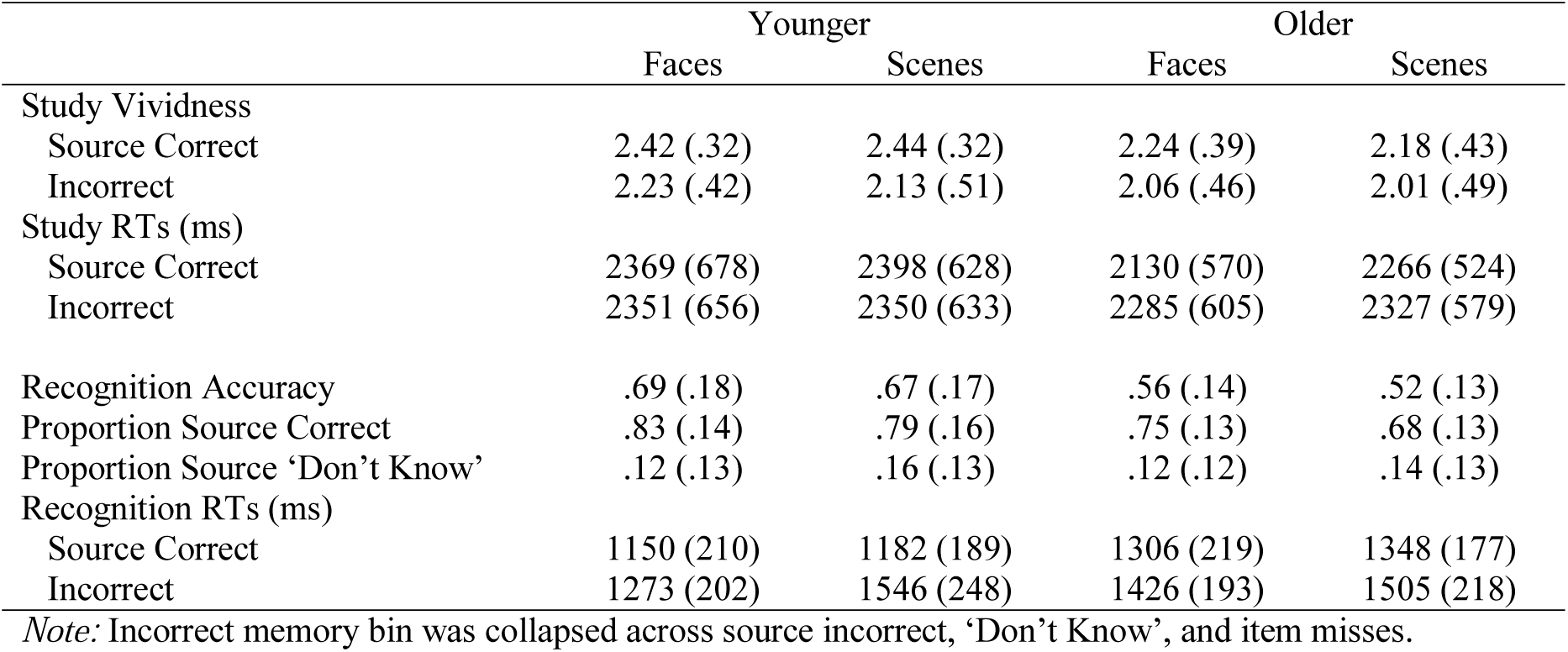
Means (SD) for behavioral performance measures.

|  | Younger |  | Older |  |
| --- | --- | --- | --- | --- |
|  | Faces | Scenes | Faces | Scenes |
| Study Vividness |  |  |  |  |
| Source Correct | 2.42 (.32) | 2.44 (.32) | 2.24 (.39) | 2.18 (.43) |
| Incorrect | 2.23 (.42) | 2.13 (.51) | 2.06 (.46) | 2.01 (.49) |
| Study RTs (ms) |  |  |  |  |
| Source Correct | 2369 (678) | 2398 (628) | 2130 (570) | 2266 (524) |
| Incorrect | 2351 (656) | 2350 (633) | 2285 (605) | 2327 (579) |
| Recognition Accuracy | .69 (.18) | .67 (.17) | .56 (.14) | .52 (.13) |
| Proportion Source Correct | .83 (.14) | .79 (.16) | .75 (.13) | .68 (.13) |
| Proportion Source ‘Don’t Know’ | .12 (.13) | .16 (.13) | .12 (.12) | .14 (.13) |
| Recognition RTs (ms) |  |  |  |  |
| Source Correct | 1150 (210) | 1182 (189) | 1306 (219) | 1348 (177) |
| Incorrect | 1273 (202) | 1546 (248) | 1426 (193) | 1505 (218) |

*Note:* Incorrect memory bin was collapsed across source incorrect, ‘Don’t Know’, and item misses.

Item recognition – operationalized as the difference between the proportion of old items correctly endorsed as ‘old’ (hit rate) and the proportion of new items erroneously endorsed as ‘old’ (false alarm rate) – was computed separately for each image category and submitted to a 2 × 2 mixed factorial ANOVA with factors of age (young, older) and stimulus category (faces, scenes). This analysis produced significant main effects of age (*F*_(1,46)_ = 10.11, *p* = .003, partial-η^2^ = .18) and category (*F*_(1,46)_ = 5.46, *p* = .024, partial-η^2^ = .11). The interaction between age and category was not significant (*F*_(1,46)_ = 0.74, *p* = .393, partial-η^2^ = .02). Post-hoc contrasts revealed that, across both categories, recognition accuracy was reduced in older relative to younger adults and that, across both age groups, accuracy was higher for items studied with faces.

Source memory accuracy was estimated using a single high-threshold model^32^ corrected for guessing^33^ using the formula *pSR* = [*pHit* - .5 * (1 - *pDK*)] / [1 - .5 * (1 - *pDK*)], where pSR refers to the probability of source recollection, and pHit and pDK refer to the proportion of correct old responses attracting an accurate or a ‘Don’t Know’ source memory endorsement, respectively. A *t*-test revealed that source memory accuracy was significantly lower in older (*M* = .51, *SD* = .16) than in younger (*M* = .68, *SD* = .18) adults (t_(45.51)_ = -3.44, p = .001). To further unpack the effect of stimulus category on source accuracy, we computed proportions of SC trials (SC/SC+SIDK) and submitted these to a 2 (age) x 2 (stimulus category) mixed-factorial ANOVA. This revealed significant main effects of age (*F*_(1,46)_ = 6.26, *p* = .016, partial-η^2^ = .12 - consistent with the foregoing analysis of the pSR metric) and category (*F*_(1,46)_ = 12.04, *p* = .001, partial-η^2^ = .21). The interaction between age and category was not significant (*F*_(1,46)_ = 0.36, *p* = .553, partial-η^2^ = .01). Post-hoc tests revealed that, across both age groups, correct source judgments were more likely for faces than scenes.

Median recognition memory RTs were sorted into correct and incorrect memory bins and submitted to a 2 (age group) x 2 (stimulus category) x 2 (memory) mixed factorial ANOVA. This gave rise to a significant three-way interaction (*F*_(1,46)_ = 8.77, *p* = .005, partial-η^2^ = .16). Post-hoc tests revealed that RTs were significantly slower for older adults relative to younger adults on face trials attracting correct (*t*_(86.8)_ = 2.60, *p* = .011) and incorrect (*t*_(86.8)_ = 2.55, *p* = .013) source judgments, as well as on correct scene trials (*t*_(86.8)_ = 2.77, *p* = .007). The two age groups did not differ on incorrect scene trials (*t*_(86.8)_ = -0.68, *p* = .501). Correct rejection RTs in young (*M* = 1384 ms, *SD* = 295 ms) and older (*M* = 1414 ms, *SD* = 213 ms) adults did not significantly differ (*t*_(41.85)_ = 0.41, *p* = .688).

### Univariate Reinstatement Effects

Trials from the study phase were binned according to category (face trials vs. scene trials) and contrasted with one another to identify patterns of encoding-related mean-signal change that were selective for each category across age groups. The results of these analyses are described in Table 2 and illustrated in Figure 1A. Face-selective encoding effects were observed in a bilateral cluster falling along the border of the cuneus and precuneus (and extending into the posterior cingulate), medial prefrontal cortex, and anterior medial temporal lobes (overlapping amygdala and anterior hippocampus and extending into right fusiform and middle temporal gyri) as well as left lateralized middle temporal gyrus and angular gyrus. Scene-selective encoding effects were evident in bilateral parahippocampal cortex (extending into retrosplenial cortex and occipital cortex), right precuneus (extending into posterior cingulate), as well as left middle and inferior frontal gyri.

**Table 2.** Loci of Univariate Encoding and Reinstatement Effects.

| Contrast | Region | MNI |  |  | Peak z | Cluster size |
| --- | --- | --- | --- | --- | --- | --- |
|  |  | x | y | z |  |  |
| Face Encoding | **Precuneus** | **3** | **-58** | **23** | **Inf** | **605** |
|  | Cuneus | 0 | -88 | 20 | 4.01 |  |
|  | WM | -18 | -46 | 17 | 3.71 |  |
|  | **Amygdala** | **21** | **-7** | **-22** | **Inf** | **2237** |
|  | Hippocampus | -24 | -7 | -22 | 7.62 |  |
|  | Middle Temporal Gyrus | 60 | -58 | 5 | 6.67 |  |
|  | **Superior Medial Gyrus** | **-3** | **59** | **17** | **6.96** | **838** |
|  | Mid Orbital Gyrus | 3 | 38 | -7 | 5.91 |  |
|  | Superior Frontal Gyrus | -12 | 44 | 44 | 3.99 |  |
|  | **Angular Gyrus** | **-57** | **-67** | **23** | **4.74** | **130** |
|  | Angular Gyrus | -48 | -64 | 26 | 4.63 |  |
|  |  | -54 | -70 | 32 | 4.60 |  |
|  | **Middle Temporal Gyrus** | **-57** | **-10** | **-16** | **4.28** | **170** |
|  | Middle Temporal Gyrus | -66 | -13 | -13 | 4.26 |  |
|  | Middle Temporal Gyrus | -54 | -31 | -7 | 4.23 |  |
| Scene Encoding | **Parahippocampal Cortex** | **30** | **-40** | **-19** | **Inf** | **6736** |
|  | Fusiform Gyrus | -27 | -46 | -16 | Inf |  |
|  | Middle Occipital Gyrus | 42 | -82 | 17 | Inf |  |
|  | **Precuneus** | **9** | **-43** | **41** | **6.84** | **324** |
|  | Precuneus | -9 | -40 | 44 | 6.66 |  |
| Face Reinstatement | **Precuneus** | **0** | **-55** | **26** | **6.31** | **151** |
|  | Precuneus | 9 | -55 | 26 | 5.33 |  |
| Scene Reinstatement | **Parahippocampal Cortex** | **27** | **-34** | **-25** | **7.31** | **405** |
|  | Retrosplenial Cortex | 15 | -55 | 11 | 5.26 |  |
|  | Parahippocampal Gyrus | 21 | -19 | -31 | 4.06 |  |
|  | **Parahippocampal Cortex** | **-24** | **-40** | **-22** | **6.92** | **224** |
|  | Fusiform Gyrus | -33 | -55 | -25 | 3.45 |  |
|  | **Retrosplenial Cortex** | **-12** | **-58** | **5** | **5.32** | **170** |

Note that for reinstatement loci, peak values were derived from the category-selective recollection contrast (inclusively masked with the corresponding encoding contrast).

**Figure 1.**
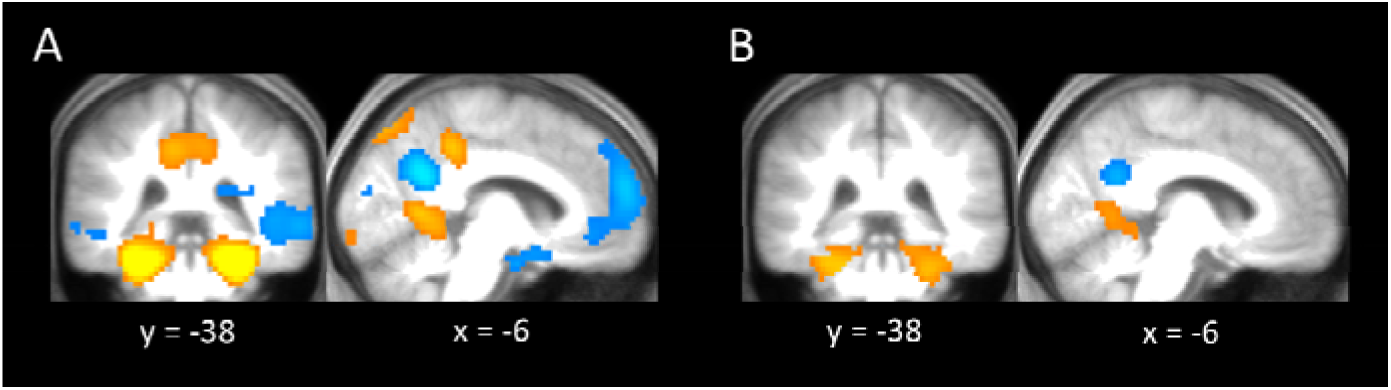
Univariate Results. (A) category-selective encoding effects operationalized as Face > Scene (cool colors) and Scene > Face (warm colors), irrespective of subsequent memory status. (B) category-selective reinstatement effects, operationalized by inclusively masking each category-selective recollection contrast with the corresponding encoding contrast. All contrasts represent main effects collapsed across age-group.

Cortical reinstatement effects were operationalized as regions where the category-selective recollection effects overlapped with the corresponding encoding effects using a sequence of masking procedures similar to those reported by Wang et al.^19^ (see Online Methods). These analyses identified face reinstatement effects in the precuneus, and scene reinstatement effects in bilateral parahippocampal cortex, extending into retrosplenial cortex. The effects are listed in Table 2 and illustrated in Fig 1B.

### Multi-Voxel Reinstatement Effects

We performed PSA to quantify cortical reinstatement effects in regions identified by the foregoing univariate analyses. PSA was performed on voxel sets identified as the top 151 voxels showing the largest z-scores within each of the category-selective reinstatement effects reported above (see Online Methods). To this end, we computed estimates of pattern similarity for each participant at two levels of specificity, namely category- and item-levels. All pattern similarity estimates were computed from single-trial *β* weights and were based on Fisher *z*-transformed Pearson’s correlation coefficients. An estimate of category-level similarity was computed to identify reinstatement of neural activity associated with each class of study context. We also computed an estimate of item-level similarity to identify reinstatement of neural activity reflecting retrieval of source information idiosyncratic to a specific study event. Due to insufficient source incorrect trial numbers, ‘incorrect’ memory bins were expanded to include ‘Don’t Know’ responses and item misses (see Online Methods). Note that unless otherwise noted, excluding the outliers identified in Figs 2 and 3 (> 3 SDs from the group means) did not alter the results reported below.

**Figure 2.**
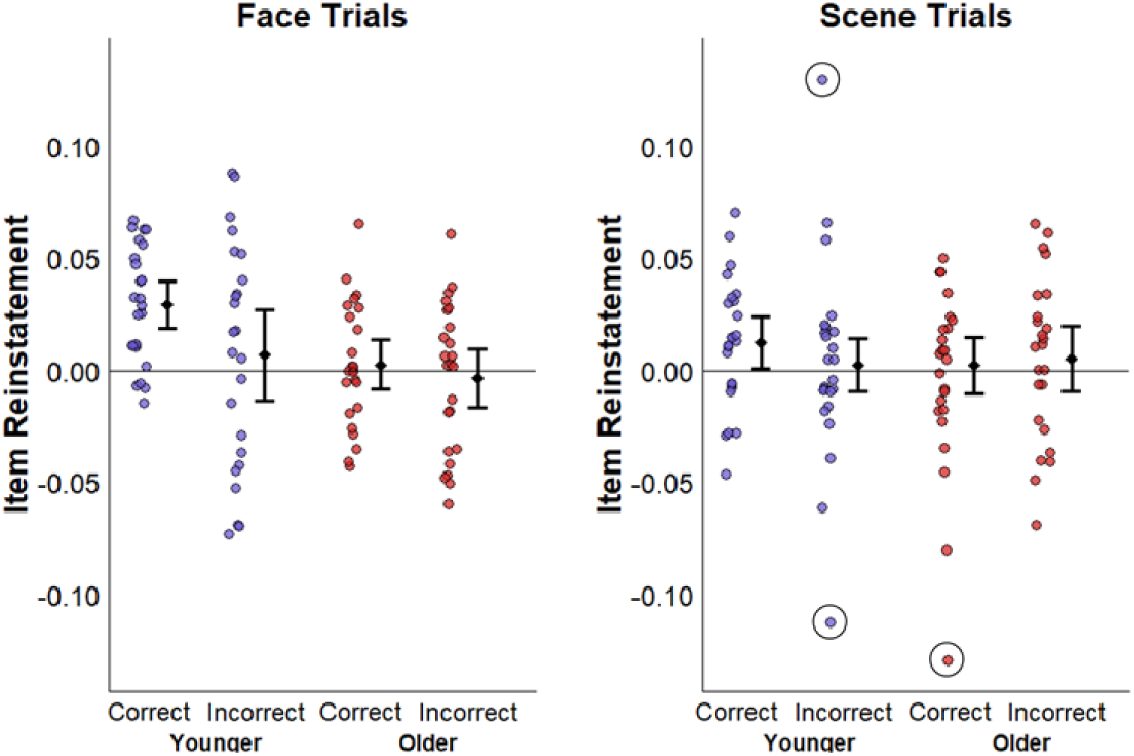
Item-level reinstatement effects. Plots illustrating across-voxel item-level pattern similarity for faces and scenes. Error bars reflect 95% confidence intervals which were computed after excluding the flagged outliers.

**Figure 3.**
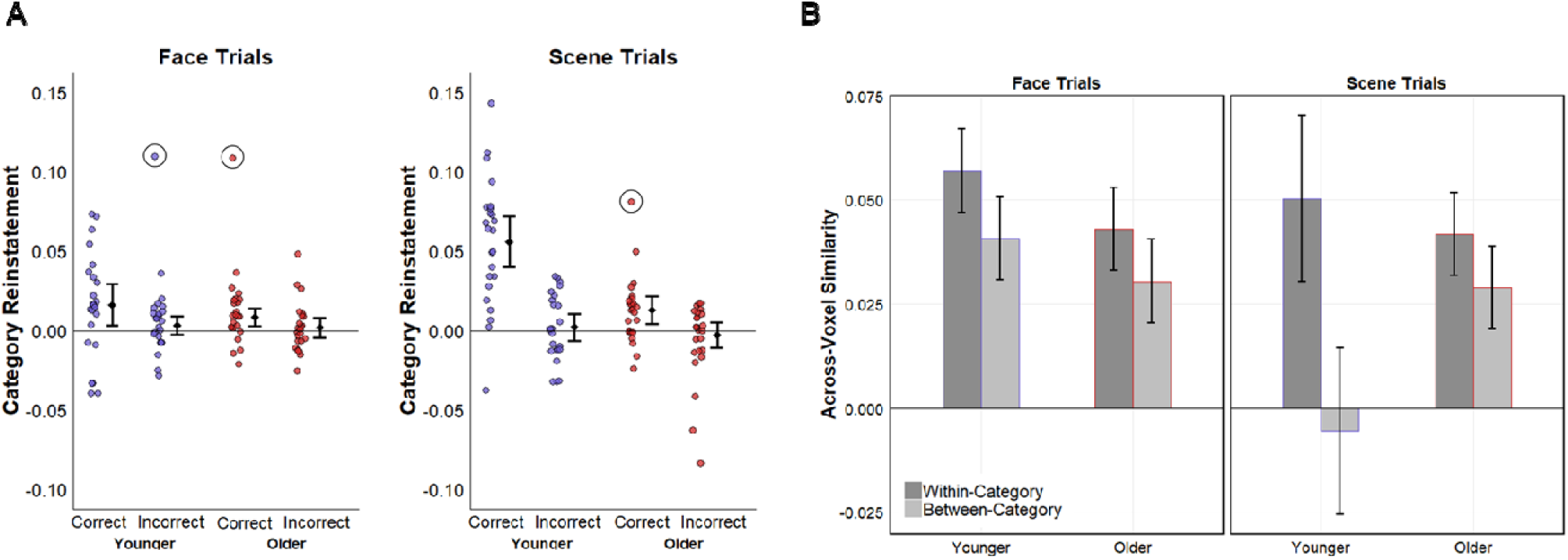
Category-Level Reinstatement Effects. (A) Plots illustrating across-voxel category-level study-test pattern similarity. Error bars reflect 95% confidence intervals, which were computed after excluding the flagged outliers. (B) Within- and between-category similarity indices are presented for each stimulus category for their respective preferred voxel set. Note that the similarity metrics shown are for source correct trials only. Error bars reflect 95% confidence intervals for the difference between mean within- and between-category pattern similarity.

To identify significant reinstatement effects, we submitted the item- and category-level PSA estimates to one-sample *t*-tests (two-tailed) against a zero null separately for each age group, image category, and memory outcome. Outcomes of the individual *t*-tests were adjusted for multiple comparisons separately for item- and category-level effects using the Holm-Bonferroni procedure. As can be seen in Fig 2, item-level pattern similarity in younger adults was significantly greater than zero for face trials attracting a correct source memory judgment (*M* = .03, *SD* = .03, *t*_(23)_ = 5.58, *p* = 1.13 × 10^−5^), and this effect remained significant after correcting for multiple comparisons. Item-level pattern similarity in younger adults did not significantly vary from zero in any of the other trial bins, and nor did it significantly differ in any of the older adult trial bins (all *ps* > .04, uncorrected).

We next submitted the item-level pattern similarity metrics to a three-way mixed-factorial ANOVA with factors of age (younger, older), category (faces, scenes), and memory status (correct, incorrect). This analysis revealed a significant main effect of age (*F*_(1,46)_ = 6.67, *p* = .010, partial-η^2^ = .13), which was driven by greater item-level pattern similarity in younger relative to older adults (*t*_(46)_ = -2.58, *p* = .013). The main effects of memory and category were not significant, and nor were any of the two- and three-way interactions (all *ps* > .1).

Turning to the category-level reinstatement effects, pattern similarity for scene trials attracting a correct source memory judgment was significantly greater than zero in younger (*M* = .06, *SD* = .04, *t*_(23)_ = 6.84, *p* = 5.59 × 10^−7^) and older (*M* = .01, *SD* = .02, *t*_(23)_ = 2.95, *p* = .007) adults, and these effects remained significant after correcting for multiple comparisons. Pattern similarity for face trials receiving a correct source memory judgment were similarly greater than zero in younger (*M* = .02, *SD* = .03, *t*_(23)_ = 2.45, *p* = .022) and older (*M* = .01, *SD* = .02, *t*_(23)_ = 2.50, *p* = .020) adults, though neither effect remained significant after correcting for multiple comparisons. When combined across age groups, however, pattern similarity for correct face trials was robustly greater than zero (*t*_(47)_ = 3.49, *p* = .001). Category-level pattern similarity effects for incorrect memory trials for both age groups and stimulus categories was far from significant (all *ps* > .15).

A three-way mixed-factorial ANOVA of category-level pattern similarity with factors of age, stimulus category, and memory status revealed a significant three-way interaction (*F*_(1,46)_ = 7.62, *p* = .008, partial-η^2^ = .14). To further unpack these results, we performed subsidiary 2 (age) x 2 (memory) mixed-factorial ANOVAs on the scene and face pattern similarity metrics respectively. The analysis of scene reinstatement effects revealed a significant age by memory interaction (*F*_(1,46)_ = 7.82, *p* = .008, partial-η^2^ = .15) which was driven by greater pattern similarity in younger relative to older adults for correct (*t*_(89.2)_ = -5.34, *p* < .0001) but not incorrect (*t*_(89.2)_ = -1.04, *p* = .300) memory trials. Additionally, scene pattern similarity was significantly greater for correct relative to incorrect memory trials in both young (*t*_(46)_ = 6.14, *p* < .0001) and older (*t*_(46)_ = 2.19, *p* = .034) adults. An analogous analysis of face pattern similarity revealed nonsignificant main effects of memory and age, as well as a nonsignificant age by memory interaction (all *ps* > .07). However, after excluding the two outlying data points evident in Fig 3, the ANOVA gave rise to a marginally significant main effect of memory (*F*_(1,44)_ = 4.05, *p* = .050, partial-η^2^ = .08).

In light of the age differences in retrieval RTs reported above, we performed a follow-up ANCOVA to contrast source correct scene reinstatement effects in young and older adults while controlling for RT. This analysis gave rise to identical results to the original ANOVA.

### Weaker scene reinstatement effects in older adults are driven by greater between-category similarity

The above-described category-level reinstatement effects were operationalized as the difference in the mean across-voxel correlation between study-test pairs belonging to the same stimulus category and the mean correlation between study-test pairs belonging to the opposite stimulus category. Therefore the influence of age on these effects is ambiguous with respect to whether the age differences were driven by reduced within-category neural similarity or increased between-category similarity. To examine this issue, the within- and between-category similarity measures elicited on source correct trials by the preferred stimulus categories for each voxel set were submitted to separate 2 (age) x 2 (similarity: within, between) mixed-factorial ANOVAs. (Note that this analysis was restricted to source correct trials as no age differences were evident on incorrect memory trials). The relevant data are presented in Fig 3B. The analysis of scene similarity measures revealed a significant interaction between age and similarity (*F*_(1,46)_ = 21.61, *p* = 2.83 × 10^−5^, partial-η^2^ = .32). Post-hoc tests revealed that the between-category similarity metric was significantly greater in older relative to younger adults (*t*_(54)_ = 2.11, *p* = .040) while the within-category similarity metric did not significantly differ between the two age groups (*t*_(54)_ = -0.53, *p* = .599). Turning to faces, there was a significant main effect of similarity (*F*_(1,46)_ = 11.99, *p* = .001, partial-η^2^ = .21) which, unsurprisingly given the data illustrated in Figure 3A, was driven by greater within-relative to between-category similarity. The main effect of age was not significant, and nor was the interaction between age and similarity type (*ps* > .1).

### Neural differentiation at encoding covaries with strength of reinstatement at retrieval

We next asked whether the above-described effects of age on category-level scene reinstatement were related to the specificity with which study events were processed at the time of encoding (i.e., neural differentiation). We computed the category-level specificity of the neural responses elicited at study by the preferred vs. the non-preferred category in each of the voxel sets employed for the analysis of reinstatement effects (see Online Methods). As before, we limited this analysis to source correct trials as it was these trials where age differences were observed. As is evident in Fig 4A, one young and one older adult had neural scene differentiation estimates that were extreme outliers (> 3 SDs from the group mean) and therefore these participants were dropped from the following analyses.

**Figure 4.**
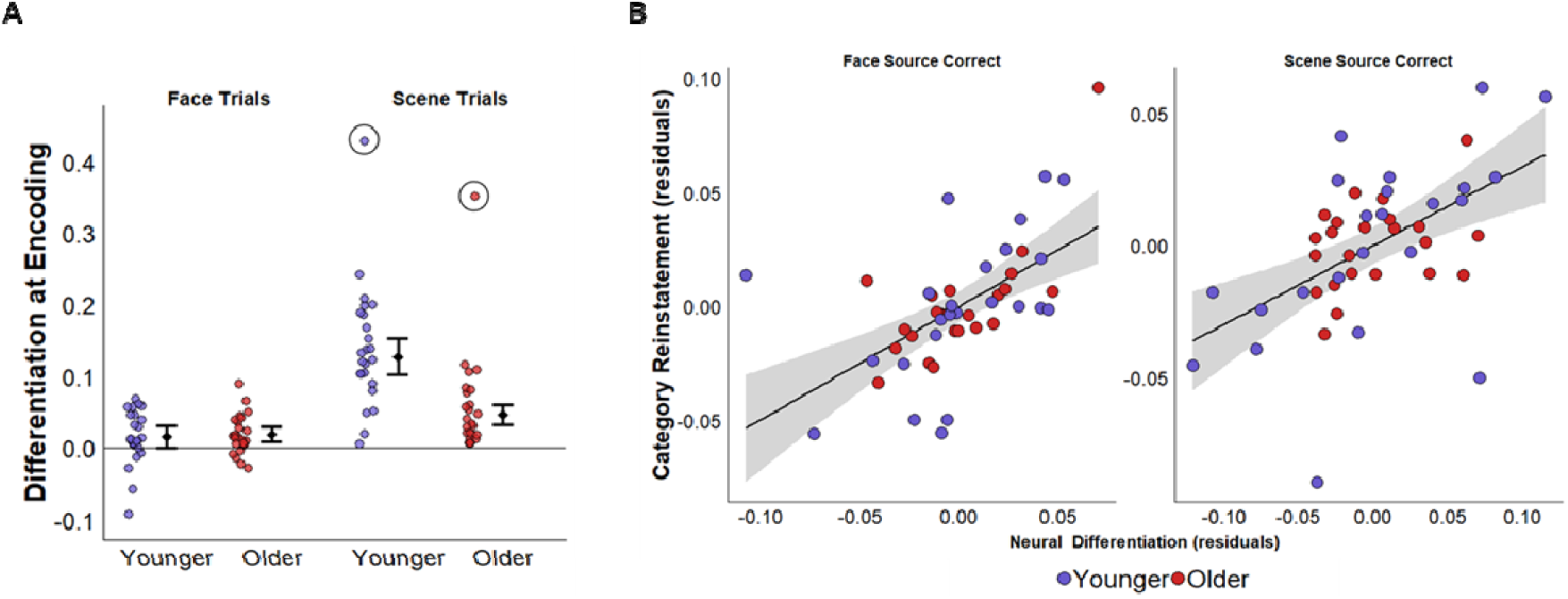
(A) Category-level similarity at encoding (i.e., neural differentiation). Error bars reflect 95% confidence intervals which were computed after excluding the flagged outliers. (B) Partial correlation between neural differentiation at encoding and category-level reinstatement at retrieval, controlling for age.

Scene pattern similarity at encoding was significantly greater than zero in younger (*M* = .13, *SD* = .06, *t*_(22)_ = 9.93, *p* = 1.37 × 10^−9^) and older (*M* = .05, *SD* = .03, *t*_(22)_ = 6.43, *p* = 1.83 × 10^−6^) adults, and these effects remained significant after correcting for multiple comparisons. Pattern similarity for face trials were similarity greater than zero in younger (*M* = .02, *SD* = .04, *t*_(23)_ = 2.07, *p* = .050) and older (*M* = .02, *SD* = .03, *t*_(23)_ = 3.46, *p* = .002) adults, though only the effect for older adults remained significant after correcting for multiple comparisons. When combined across age groups, however, pattern similarity for face trials was robustly greater than zero (*t*_(47)_ = 3.73, *p* = .001). A 2 (age) x 2 (category) factorial ANOVA revealed a significant age group x category interaction (*F*_(1,46)_ = 10.97, *p* = .002, partial-η^2^ = .19). Neural differentiation was significantly lower for scenes in older relative to younger adults (*t*_(91.7)_ = -4.63, *p* < .0001) but, echoing the category-level reinstatement effects illustrated in Fig 3A, no age differences were observed for face differentiation (*t*_(91.7)_ = 0.18, *p* = .855).

For each stimulus category, we computed the partial correlation between neural differentiation at encoding and category-level reinstatement, controlling for age. The resulting partial correlations were highly significant for both faces (*r*_*partial*_ = .58, *p* = 2.26 × 10^−5^) and scenes (*r*_*partial*_ = .53, *p* = 1.63 × 10^−4^) and multiple regression analyses indicated that age did not significantly moderate either of these relationships (*ps* > .2 for both interaction coefficients). In light of these findings, we conducted an ANCOVA to contrast category-level scene reinstatement in young and older adults while controlling for neural differentiation at study. The analysis revealed a non-significant age effect (*F*_(1,43)_ = 3.82, *p* = .057, partial-η^2^= .08). When the analysis was conducted after excluding the covariate, the main effect of age was of course highly significant (*F*_(1,44)_ = 26.11, *p* = 6.71 × 10^−6^), and the effect size increased substantially (η^2^= .37). In short, the inclusion of neural differentiation at study as a covariate led to a nearly five-fold decrease in the proportion of variance in scene reinstatement explained by the factor of age group.

### Retrieval-related reinstatement and hippocampal retrieval effects independently covary with source accuracy

We performed a series of generalized linear mixed-effects analyses to examine the relationships between trial-wise estimates of retrieval-related hippocampal activity, cortical reinstatement, and source memory accuracy (see Online Methods). We elected to run separate models for scene and face trials to ease interpretation and to avoid model overfitting. In each model, trial-wise binary source memory outcomes (correct, incorrect) were entered as the dependent variable. The outcomes of these analyses are reported in Table 3 and illustrated in Fig 5 below. A companion analysis of point-biserial partial correlation coefficients (see Online Methods and supplemental materials) led to conclusions closely similar to those derived from the multilevel regression models reported below.

**Table 3.**
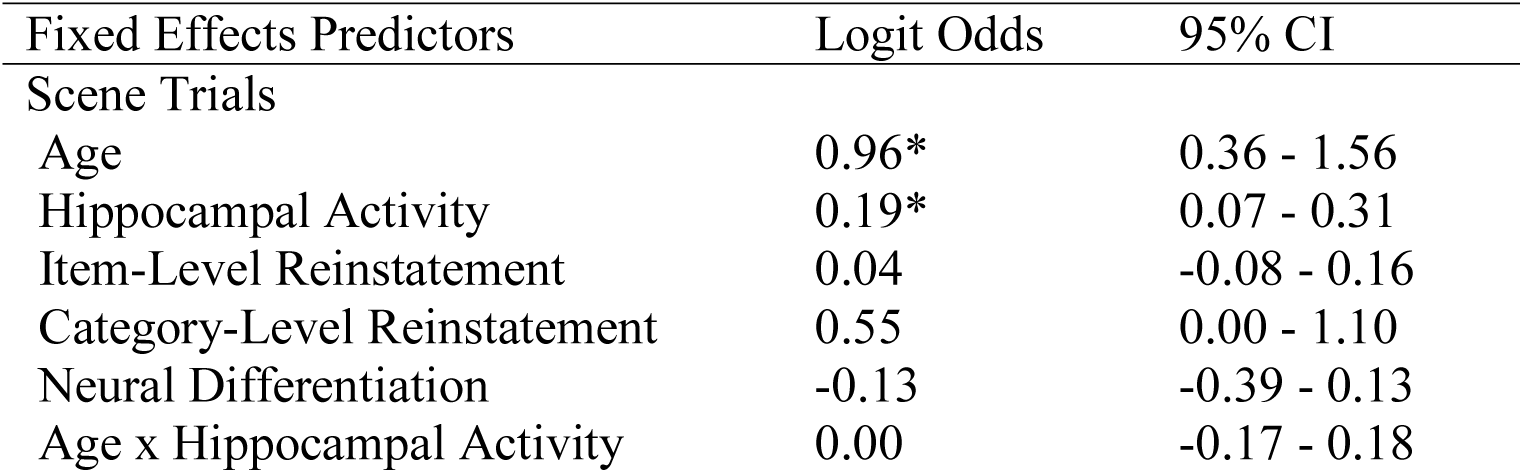
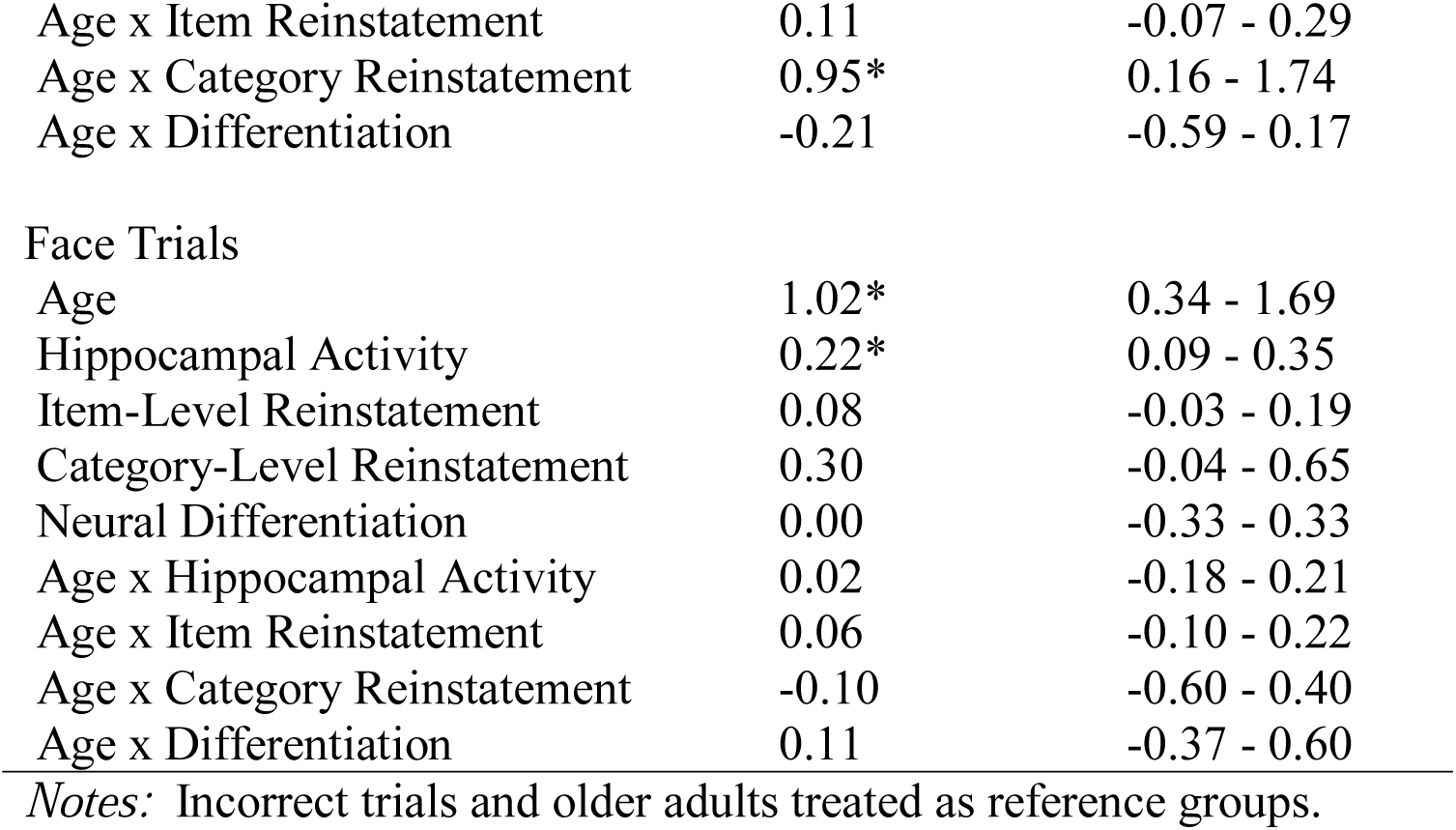
Results of generalized linear mixed effects analyses.

| Fixed Effects Predictors | Logit Odds | 95% CI |
| --- | --- | --- |
| Scene Trials |  |  |
| Age | 0.96* | 0.36 - 1.56 |
| Hippocampal Activity | 0.19* | 0.07 - 0.31 |
| Item-Level Reinstatement | 0.04 | -0.08 - 0.16 |
| Category-Level Reinstatement | 0.55 | 0.00 - 1.10 |
| Neural Differentiation | -0.13 | -0.39 - 0.13 |
| Age x Hippocampal Activity | 0.00 | -0.17 - 0.18 |
| Age x Item Reinstatement | 0.11 | -0.07 - 0.29 |
| Age x Category Reinstatement | 0.95* | 0.16 - 1.74 |
| Age x Differentiation | -0.21 | -0.59 - 0.17 |
| Face Trials |  |  |
| Age | 1.02* | 0.34 - 1.69 |
| Hippocampal Activity | 0.22* | 0.09 - 0.35 |
| Item-Level Reinstatement | 0.08 | -0.03 - 0.19 |
| Category-Level Reinstatement | 0.30 | -0.04 - 0.65 |
| Neural Differentiation | 0.00 | -0.33 - 0.33 |
| Age x Hippocampal Activity | 0.02 | -0.18 - 0.21 |
| Age x Item Reinstatement | 0.06 | -0.10 - 0.22 |
| Age x Category Reinstatement | -0.10 | -0.60 - 0.40 |
| Age x Differentiation | 0.11 | -0.37 - 0.60 |

*Notes:* Incorrect trials and older adults treated as reference groups.

**Figure 5.**
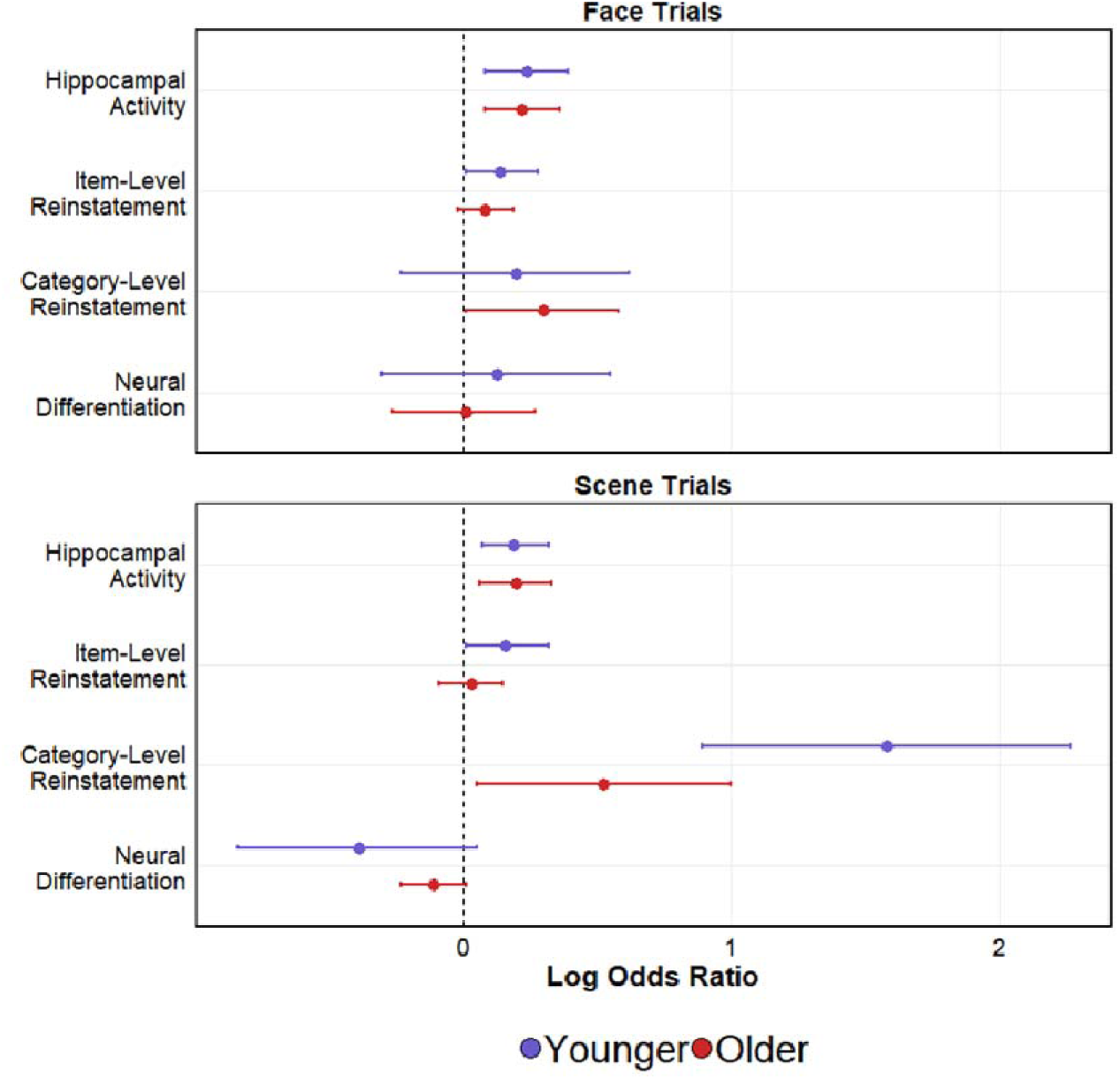
Logit odds with 95% confidence intervals are plotted for each fixed effects predictor, separated by age.

As is evident from Table 3 and Fig 5, the odds of making a correct source memory judgment significantly co-varied with increasing hippocampal activity for both face and scene trials. For face trials, neither reinstatement metric predicted memory accuracy. For scene trials, there was a significant age x category-level reinstatement interaction which was driven by a stronger within-subject relationship between reinstatement of category-level scene information and source memory performance in younger adults. As is evident in Fig 5, however, this relationship was reliably greater than chance in both age groups.

### Retrieval-related hippocampal activity covaries with reinstatement of trial-unique scene information

In a final analysis, we used linear mixed effects analyses to examine relationships between trial-wise estimates of retrieval-related hippocampal activity and reinstatement metrics. As previously, these analyses were performed separately for each stimulus category. In each model, trial-wise estimates of item- or category-level reinstatement were entered as the dependent variable. Age, hippocampal activity, and the age x hippocampal activity interaction term were entered as fixed effects predictors, and subject-wise intercept and slope terms were entered into the models as random-effects factors. The analyses revealed that hippocampal activity was a significant predictor of item-level reinstatement of scene information (*b* = .12, *p* = 2.23 × 10^−4^). The age x hippocampal interaction was not significant, nor did hippocampal activity significantly covary with category-level scene reinstatement (*b* = .01, *p* = .836). Hippocampal activity did not significantly covary with either item- (*b* = -.05, *p* = .119) or category-level (*b* =.04, *p* = .145) reinstatement of face information.

## Discussion

We examined whether the influence of age on retrieval-related cortical reinstatement was moderated by age differences in neural selectivity during encoding, whether reinstatement covaried with source memory performance, and whether any such co-variation might explain age-related variance in source memory performance. Univariate reinstatement effects for scene information were identified in bilateral parahippocampal and retrosplenial cortex, canonical scene selective regions^34^. Univariate reinstatement effects for face information were evident in bilateral precuneus, a prominent member of the ‘extended’ face processing network^35^. Using multi-voxel PSA in ROIs defined by the univariate analyses, we identified robust age differences in the strength of recollection-related reinstatement of scene information which were fully explained by analogous differences in neural differentiation at the time of encoding. In addition, there was a significant relationship between trial-wise metrics of scene reinstatement and memory accuracy in both age groups, although the relationship was significantly stronger in the younger group.

Turning first to the behavioral results, consistent with prior reports^36^ vividness ratings at study predicted memory performance, but this relationship did not vary with age. The age differences we observed in neural differentiation and reinstatement are therefore unlikely to reflect the confounding effects of this variable. At test, source memory performance was lower in older adults (and was accompanied by an analogous effect for item memory), as would be expected given the extensive prior literature documenting age-related episodic memory decline (for review, see^3^). Regardless of age, both item and source memory were higher for test words paired with faces than with scenes. While these memory benefits for faces are not without precedent^37,38^ (but see^30,39^) they currently lack an explanation. Since the effects did not interact with age group, we do not discuss them further here.

We used multivoxel PSA to examine whether face and scene reinstatement effects associated with successful source memory judgments differed in strength according to age. Category-level reinstatement of scene information was robustly weaker in older adults relative to younger individuals, a finding consistent with some prior reports^12,13,15^ but inconsistent with others^18,19,40^. This age-related attenuation in neural specificity was driven not by a reduction in levels of within category (scene-scene) similarity, but rather, by elevated between category (scene-face) similarity. This finding is arguably analogous to the ‘neural broadening’ effects which have been reported as a possible source of age-related neural dedifferentiation in studies employing univariate analysis methods^41^ (but see^24^ for an example of age-related dedifferentiation driven by neural attenuation).

In line with prior findings (e.g.^24,29^), we identified age differences in scene differentiation not only at retrieval, but at encoding also. Echoing the findings for retrieval, scene selectivity in the same voxel set was robustly lower in the older group at encoding. These and similar findings have been interpreted as evidence for age-related neural dedifferentiation, a reduction in neural specificity with age that has been proposed to contribute to age-related cognitive decline (for review, see^22^). (We note that the present findings of neural age-related dedifferentiation did not extend to faces; this result echoes prior observations for visual objects, and will be discussed in a separate paper). Crucially, the age differences that we identified in recollection-related reinstatement effects in the scene voxel set were eliminated when the scene similarity metric derived from the same voxel set at encoding was employed as a covariate. Thus, the present findings suggest that age differences in retrieval-related reinstatement are fully attributable to the selectivity with which the retrieved information was represented in category-selective cortex when it was initially experienced. In other words, the present findings offer scant support for proposals that age-related memory decline reflect an impairment in retrieval processes that support recollection of detailed information (e.g.^12,15^). Also of importance, across both stimulus categories the relationship between neural differentiation at encoding and the strength of retrieval-related reinstatement was age-invariant (Figure 4B), suggesting that the relationship identified here between encoding- and retrieval-related neural specificity reflects a general principle of brain function that operates across the adult lifespan.

In addition to the aforementioned scene reinstatement effects, we also identified reliable category-level reinstatement of face information in both young and older adults. Echoing the findings from the encoding phase, there was no evidence that these face reinstatement effects differed with age. These findings are arguably consistent with the proposal advanced above that age differences in retrieval-related reinstatement depend on the existence of analogous differences in neural differentiation at the time of encoding.

Across both stimulus categories, reinstatement of item-level information was reduced in older adults relative to their younger counterparts. However, this effect did not interact with source memory accuracy and therefore is hard to interpret in terms of its implications for the age differences in memory for the items. The weak effects for item-level reinstatement might reflect our use of a memory test that did not require retrieval of trial-unique information: perfect performance on the test would have been possible simply by recalling which of the stimulus categories a recognized test word had been associated with at study. Regardless of the validity of this account, the present findings offer little support for the proposal^19^ that age differences in cortical reinstatement effects might be more prominent when the effects are examined at the item-rather than the category-level.

Turning to the trial-wise analyses, consistent with prior findings^30^ we identified a robust within-subject relationship between category-level scene reinstatement and memory accuracy in younger adults. Although it was reliably greater than chance, the strength of this relationship was significantly attenuated among older participants (Figure 5). Although it comes from a qualitatively different analysis approach to that employed in prior aging studies (within-rather than between-participants correlations between neural and behavioral measures), this finding marks a departure from prior observations that the relationship between neural differentiation and memory performance is not strongly moderated by age (for review, see^22^). The age dependency of this relationship suggests that the fidelity of reinstated neural information plays a less important role in mediating memory success in older than in young adults.

Consistent with the prior findings of Gordon et al.^30^ and Richey et al.^31^, trial-wise estimates of retrieval-related hippocampal activity covaried within-subjects with source memory accuracy for both faces and scenes. Of importance, these effects were invariant with respect to age, consistent with prior proposals that recollection-related hippocampal activity does not differ in magnitude with age and demonstrates an age-invariant relationship with memory performance (at least for cognitively unimpaired older adults)^19,42^. On their face, the present hippocampal retrieval effects are consistent with the widely accepted notion that successful episodic retrieval depends on the hippocampal-mediated ‘reactivation’ of patterns of cortical activity encoded in the hippocampus as an episode was initially experienced (e.g.^10,43,44^. Nevertheless, for scene trials, hippocampal activity and the strength of category-level reinstatement independently predicted memory performance. Moreover, retrieval-related hippocampal activity did not explain a significant fraction of the variance in category-level scene reinstatement. Thus, we found no evidence that cortical reinstatement of scene information mediated (even partially) the relationship between retrieval-related hippocampal activity and memory performance (cf.^30,31^). The reasons for this disparity between the present and prior findings are unclear. While it is possible that we may have identified a relationship with reinstatement had we employed more nuanced measures of retrieval-related hippocampal activity (e.g. at the level of individual cell fields) we note that Gordon et al.^30^ employed a very similar approach to that used here to define a whole hippocampus ROI.

In contrast to the foregoing results, we did identify a robust age-invariant relationship between hippocampal activity and reinstatement of trial-unique scene information. One possibility is that retrieval-related hippocampal activity is preferentially involved with reactivation of trial-unique information, rather than reinstatement of category-level patterns of activity^31,45^ (but see also^30^). This possibility is undermined, however, by our failure to identify any evidence that item-level reinstatement of scene (or face) information covaried with memory performance. This result should therefore be interpreted with caution.

In conclusion, we identified robust age differences in retrieval-related scene reinstatement which could be explained by analogous differences in neural differentiation at encoding. Importantly, for both faces and scenes, the relationship between neural differentiation at encoding and strength of retrieval-related reinstatement did not differ with age. These findings suggest that, regardless of age, the specificity with which events are neurally processed at the time of encoding determines the fidelity of cortical reinstatement at retrieval.

## Methods

### Participants

Participants were 27 younger and 33 older adult volunteers recruited from the University of Texas at Dallas and surrounding community. All participants were right-handed and fluent English speakers before the age of five. No participants had a history of neurological or psychiatric disease or reported taking any prescription medications affecting the central nervous system. All participants gave informed consent in accordance with the UT Dallas and University of Texas Southwestern Institutional Review Boards and were compensated at the rate of $30 an hour.

Data from three younger and three older adult participants were excluded from subsequent analyses for the following reasons: voluntary withdrawal (n=2), behavioral performance resulting in memory bins with too few trials (n=2), technical malfunction (n=1), and an incidental MRI abnormality (n=1). An additional six older adult participants were excluded due to near-chance source memory performance according to a pre-determined cut-off score (pSR < .1). Data from the remaining 24 younger (18-28 years, *M* = 22.4 years, *SD* = 3.2 yrs; 15 females) and 24 older (65-75 years, *M* = 70.1 yrs, *SD* = 3.4 yrs; 14 females) participants were used in the analyses reported here.

### Neuropsychological Testing

All participants completed a neuropsychological test battery consisting of the Mini-Mental State Examination (MMSE), the California Verbal Learning Test-II^46^ (CVLT), Wechsler Logical Memory Tests 1 and 2^47^, Trail Making tests A and B^48^, the Symbol Digit Modalities test^49^ (SDMT), the F-A-S subtest of the Neurosensory Center Comprehensive Evaluation for Aphasia^50^, the Wechsler Adult Intelligence Scale–Revised subtests of forward and backward digit span^51^, category fluency^52^, Raven’s Progressive Matrices^53^ (List 1) and the Wechsler Test of Adult Reading^51^ (WTAR). Participants were excluded from entry into the study if they scored < 27 on the MMSE, < 1.5 SDs below age-appropriate norms on any memory test, < 1.5 SDs below age norms on any two other tests, or if their estimated full-scale IQ was less than 100. These criteria were employed to minimize the likelihood of including individuals with mild cognitive impairment. Results from the neuropsychological test battery are presented in supplementary Table 1.

### Experimental Materials and Procedure

The critical experimental stimuli consisted of 288 concrete words and 96 color images depicting faces (50% male) and a further 96 images depicting scenes (50% urban, 50% rural). An additional 68 words and 40 images were used as filler or practice stimuli.

Separate stimulus sets were created for yoked younger-older adult pairs. For the study phase, each stimulus set comprised 192 randomly selected critical image-word pairs interspersed with 96 null trials (white fixation cross). Study sets were divided evenly into four sub-lists (one per scanning session) of 48 critical and 24 null trials. Stimulus sets for the test phase comprised the 192 ‘old’ words encountered during the study task interspersed with 96 unstudied ‘new’ words and 96 null trials. Test stimuli were subdivided evenly into four sub-lists (one per scanning session), each comprising 48 ‘old’ items, 24 ‘new’ items, and 24 null trials. Trial order for both tasks was pseudo-randomized under the constraint that no more than three critical trials from the same encoding category (faces, scenes) or two null trials occurred consecutively.

The experiment consisted of two study-test cycles completed inside the scanner. The study and test tasks were split into two scan sessions each per cycle. The sequence and timing for each trial was identical in the study and test phases. Each trial commenced with a red fixation cross (500 ms) followed by the presentation of the study (word-image pair) or test (word only) items for 2000 ms. Each item was followed by a white fixation cross displayed for an additional 2000 ms, resulting in a stimulus-onset-asynchrony (excluding null trials) of 4.5 s. A 30 s rest break occurred midway through each scan session. Two filler trials were presented at the beginning of each scan session and after each rest break. Instructions and practice for the study and test tasks were given outside of the scanner.

For each study trial, participants viewed a concrete noun paired with an image of a face or a scene. All face and scene images were scaled and cropped to 256 × 256 pixels. On face trials, participants were instructed to imagine the person depicted in the image interacting with or using the object denoted by the word. On scene trials, participants were instructed to imagine the object denoted by the word interacting or moving about within the depicted scene. To encourage compliance with the instructions, participants rated the vividness of the imagined scenario on a 3-point scale (‘Not Very Vivid’, ‘Somewhat Vivid’, ‘Very Vivid’). Responses were made on a scanner-compatible button box with the index, middle, and ring fingers of the right hand. Responses made within 500-4500 ms of the presentation of the study items were included in the analyses. Trials associated with responses falling outside this temporal window were discarded and included as events of no interest.

During the test phase, participants viewed old and new words presented one at a time. Each test item required either one or two memory judgments. Participants were first required to judge whether a test item was previously encountered during the study phase by making an “Old” or “New” response. They were instructed to refrain from guessing and to respond “Old” only when confident that a test item had been previously studied. For items attracting an “Old” response, participants were then required to make a source memory judgment concerning the associated encoding category (i.e., was the word studied with a face or a scene?). A “Don’t Know” response option was also available to discourage guessing. Responses were made on a scanner compatible button box with the index, middle and ring fingers of the right hand. Old/new recognition judgments were always made with the index and middle fingers and counterbalanced across participants. Source memory responses were also fully counterbalanced across participants with the constraint that the “Don’t Know” response was never assigned to the middle finger. As with the study phase, trials associated with responses made between 500ms and 4500 ms of the presentation of the test items were included in the analyses. Trials falling outside this temporal window were discarded and included as events of no interest in the design matrix.

### MRI Data Acquisition and Preprocessing

Functional and anatomical images were acquired with a 3T Philips Achieva MRI scanner (Philips Medical Systems, Andover, MA, USA) equipped with a 32-channel receiver head coil. Functional images were acquired using a T2*-weighted, blood-oxygen level-dependent echoplanar (EPI) sequence (sensitivity encoding [SENSE] factor 2, flip angle 70 deg., 80 × 78 matrix, field of view [FOV] = 24 cm, repetition time [TR] = 2000 ms, and echo time [TE] = 30 ms). EPI volumes consisted of 34 slices (1-mm interslice gap) with a voxel size of 3×3×3 mm. Slices were acquired in ascending order oriented parallel to the anterior commissure-posterior commissure plane. Each functional run included 194 (study phase) or 248 (test phase) EPI volumes. T1-weighted anatomical images were acquired with a magnetization-prepared rapid gradient echo (MPRAGE) pulse sequence (FOV = 240 × 240, 1 × 1 × 1 mm isotropic voxels, 34 slices, sagittal acquisition.

fMRI preprocessing and analyses were conducted using a combination of Statistical Parametric Mapping (SPM12, Wellcome Department of Cognitive Neurology, London, UK) and custom scripts, run under Matlab R2017a (MathWorks). Functional images were realigned to the mean EPI image and slice-time corrected using sinc interpolation to the 17^th^ slice. The images were then reoriented and spatially normalized to a sample-specific EPI template following previously published procedures^42,54^. Normalized volumes were resampled into 3 mm isotropic voxels and smoothed with an isotropic 8 mm full-width half-maximum Gaussian kernel. Anatomical images were spatially normalized to a sample-specific T1 template following procedures analogous to those applied to the functional images.

### MRI Data Analysis

fMRI data were analyzed using both mass univariate analysis and MVPA to, respectively, localize and quantitate reinstatement effects. The univariate analyses were performed in two stages. In the first stage, separate GLMs were constructed for each participant. Parameter estimates from events of interest were then carried forward to second-level random effects factorial ANOVAs to test for group level effects. Separate GLMs were employed to analyze the study and test phases.

Study trials were binned according to encoding category (i.e., face trials vs. scene trials), giving two events of interest. These were modeled with a 2 s duration boxcar regressor convolved with SPM’s canonical hemodynamic response function (HRF) and an orthogonalized delayed HRF^55^ generated by shifting the orthogonalized canonical HRF one TR (2 s) later in time. The results obtained for the late HRF added little of theoretical significance to the findings obtained with the canonical function and are not reported here. Filler trials, trials with missing or multiple responses, and trials falling outside of the aforementioned temporal response window were modeled as covariates of no interest, along with the 30 s rest periods and six regressors representing motion-related variance (three for rigid-body-translation and three for rotation). Data from volumes showing transient displacement > 1 mm or rotation > 1° in any direction were censored by their inclusion as additional covariates of no interest. Parameter estimates from the two events of interest were carried forward to a second-level random effects 2 × 2 factorial ANOVA treating age (younger, older) as a between subjects factor and category (faces, scenes) as a within subjects factor.

For the test phase, five events of interest were included in the design matrix: correct source memory for faces (SC_faces_), correct source memory for scenes (SC_scenes_), successfully recognized items eliciting an incorrect source memory endorsement (including ‘DK’ responses; SIDK), old items erroneously endorsed as new (item misses), and new items correctly endorsed as new (correct rejections). To ensure an adequate number of trials per memory bin, source incorrect and item misses were collapsed across the two encoding categories. Each event of interest was modeled with a delta function convolved with SPM’s canonical HRF and an orthogonalized delayed HRF. As before, the results obtained for the late HRF added little of theoretical significance to the findings obtained with the canonical function and are not reported here. As with the study phase, filler trials, trials with missing or multiple responses, and trials falling outside of the aforementioned temporal response window were modeled as covariates of no interest, along with the 30s rest periods and six regressors representing motion-related variance (three for rigid-body translation and three for rotation). Data from volumes showing a transient displacement of > 1 mm or 1° in any direction were eliminated by their inclusion as covariates of no interest. Parameter estimates from the five events of interest were carried over to a second-level random effects 2 × 5 factorial ANOVA treating age (younger, older) as a between subjects factor and subsequent memory (source correct faces, source correct scenes, source incorrect, item miss, correct rejection) as a within subjects factor.

### Univariate Reinstatement Effects

Using univariate analyses of mean-signal change, cortical reinstatement effects were operationalized as regions where category-selective encoding and recollection effects overlapped. The regions were identified using a sequence of masking procedures similar to those reported by Wang et al.^19^. Recollection effects were operationalized by separately contrasting source correct trials from each image category with source incorrect trials pooled across the two encoding conditions (SC_faces_ > SIDK, SC_scenes_ > SIDK; height threshold p < .001). We then exclusively masked the two recollection contrasts with one another in order to identify recollection effects that were unique to each category (mask threshold *p* < .05). The resulting category-selective recollection contrasts were then inclusively masked with the corresponding category-selective encoding contrast (Faces > Scenes or Scenes > Faces, inclusive mask threshold *p* < .001). Clusters that survived family-wise error (FWE) corrected extent thresholds in the masked recollection contrasts were retained.

### MVPA Feature Selection

Pattern similarity analysis (PSA) was performed on the 151 voxels showing the largest z-scores for each of the two category-selective reinstatement effects identified by the mass univariate analyses described above. This set size was selected to coincide with the spatial extent of the smaller of the two mass univariate reinstatement effects (Table 2). Thus, for both stimulus categories, PSA was only performed on voxels that demonstrated reliable suprathreshold reinstatement effects at our pre-experimental statistical threshold. To avoid bias, these voxel sets were empirically defined at the single participant level by using an iterative ‘leave-one-out’ approach^19^. For each of 24 randomly yoked young-older adult pairs, the top 151 voxels for each stimulus category were determined from the remaining 46 participants and assigned to the left-out pair. This ensured that the data used to define the voxel sets for each participant were independent of the data subjected to PSA.

### Pattern Similarity Analyses

Unsmoothed data from the four study sessions were concatenated and subjected to a ‘least-squares-all’ GLM^56,57^ to estimate the BOLD response for each trial separately. Each study event was modeled as a 2 s duration boxcar convolved with the canonical HRF. Unsmoothed data from the four test sessions were analyzed in a similar manner with the exception that test events were modeled with a delta function convolved with the canonical HRF. Pattern similarity analyses (PSA) were performed on the resulting single-trial *β* weights and were based on Fisher *z*-transformed Pearson correlation coefficients. A pattern similarity metric was computed separately for each image category within the respective category-selective 151-voxel sets.

Category-level reinstatement effects were defined as the difference between the mean across-voxel correlation between a given study trial and all test trials involving the same image type, and the mean correlation between that same study trial and all test trials involving the alternate image type (i.e., a within-between *category* similarity metric). This procedure was performed separately for correct (SC) and incorrect (SIDK + item miss) trials. A summary measure of category-level pattern similarity was computed for each participant by averaging across all of the trial-wise within-between similarity estimates for a given image category and source memory outcome.

Item-level reinstatement effects were defined as the difference between the mean across-voxel correlation between a given study trial and its corresponding test trial, and the mean correlation between that same study trial and all other test trials involving the same image type (giving an *item-wise* similarity metric). As was just described, item-level pattern similarity was computed separately for correct and incorrect memory trials, and a summary measure was computed for each participant as the average trial-wise within-between similarity estimates corresponding to a given image category and source memory outcome.

In addition to assessing encoding-retrieval overlap, we also employed PSA to quantify encoding-related neural differentiation in the face and scene voxel sets used to identify category- and item-level reinstatement effects (e.g.^24^). For the face-selective voxel set, the within-category measure was the average across-voxel similarity between a given face trial and all other face trials, and the between-category measure was the average correlation between a given face trial and all scene trials. The same approach was used for the scene-selective voxel set, except that scene trials were used for the within-category measures, and face trials were used for the between-category measures. A measure of neural differentiation for each voxel set was computed as the difference between the respective within and between similarity metrics, averaged across all trials. Note that PSA was computed across different scanning sessions to avoid the possibility of bias from temporal autocorrelations between trials occurring in the same scanning session^56^.

### Trial-wise Mixed Effects Analyses

Bilateral hippocampal masks were manually traced on an anatomical T1 template derived from a large cross-sectional dataset from our lab (36 younger, 36 middle-aged, and 64 older adults^58^. For each participant, we generated two unilateral vectors comprising single trial *β* weights averaged across all voxels falling within left and right hippocampal masks, respectively. Each vector was then *z-*transformed across trials separately for each participant, and the correlation (Fisher-z transformed) between left and right trial-wise hippocampal activity was computed. The mean across-participant correlation between left and right hippocampal activity was highly significant (mean *r* = .67, *p* = 2.20 × 10^−16^). Motivated by this result, and lacking any a priori hypotheses regarding hippocampal lateralization, we computed bilateral trial-wise hippocampal activity by averaging the parameter weights across the two hemispheres.

To examine the link between trial-wise estimates of retrieval-related hippocampal activity, cortical reinstatement, and source memory accuracy, we performed a set of generalized linear mixed-effects models separately for each stimulus category. In each model, trial-wise binary source memory outcomes (correct, incorrect) were entered as the dependent variable. Age, hippocampal activity, category- and item-level reinstatement, neural differentiation at encoding, and all two-way interaction terms involving age were entered as fixed effects predictors. Subject-wise intercept and slope terms were entered into the model as random-effects factors. Models were fit using maximum likelihood Laplace approximation.

In a companion analysis, we computed point-biserial correlations for each individual participant between trial-wise binary source memory judgments (SC, SIDK + item misses) and each of the four fixed effects predictors (hippocampal activity, category- and item-level reinstatement, encoding-related neural differentiation), in each case controlling for the effects of the other predictors. This procedure was performed separately for face and scene trials. The resulting correlation coefficients were Fisher-z transformed and then submitted to random effect analyses separately for each image type and fixed effect factor (see supplemental materials). The results from this procedure closely approximated those obtained from the generalized linear mixed-effects models (Fig. S1).

### Statistical Analyses

All statistical analyses were conducted with R software (R Core Team, 2017). All *t*-tests were two-tailed and performed using the t.test function in the base R package. Welch’s unequal variance *t*-tests were performed when assumptions of equal variance were not met. ANOVAs were conducted using the *afex* package^59^ and the Greenhouse-Geisser procedure^60^ was used to correct degrees of freedom for non-sphericity when necessary. Post-hoc tests on significant effects from the ANOVAs were conducted using the *emmeans* package^61^ and corrected for multiple comparisons using the Holm-Bonferroni procedure where appropriate.

Generalized linear mixed-effects models were performed using the *glmer* function in the lme4 package^62^. Point-biserial correlations were computed using the *ltm* package^63^.

## Author Contributions

M.D.R. conceived of the project. D.R.K & M.D.R designed the experiment. P.F.H performed the experiment and analyzed the data. P.F.H & M.D.R interpreted the data and wrote the manuscript.

