## Supplemental Material for "Age differences in retrieval-related reinstatement reflect age-related dedifferentiation at encoding"

**Supplemental Materials**

**Generic Univariate Recollection Effects**. To identify recollection effects that were common to the two image categories, we inclusively masked the across group main effect of the source recollection contrast (one-sided t-contrast, *p* < .001, uncorrected) with the two directional category-specific recollection contrasts (inclusive mask threshold *p* < .001). This contrast revealed three significant clusters of recollection related activity in bilateral medial temporal lobes and basal ganglia (Fig. 2C). A large cluster in the left hemisphere (xyz = -21 -7 -22, peak z = 5.65, k = 501) was centered in the hippocampus, extending into the amygdala, putamen, and nucleus accumbens. Two right lateralized clusters were located in the hippocampus (xyz = 39 -16 -22, peak z = 5.31, k = 126) and putamen (xyz = 30 -13 -1, peak z = 5.76, k = 96, significant after small volume correction). A whole-brain age x recollection interaction contrast (height-threshold *p* < .001, uncorrected) failed to identify any clusters where generic recollection effects differed according to age group.

**Across-participant item- and category-level reinstatement metrics do not predict source memory accuracy independently of age.** We ran a series of multiple regression analyses in which age group and, respectively, participant-wise item- and category-level pattern similarity metrics for each stimulus category were employed as predictors of source accuracy (pSR). All age x reinstatement interaction terms were all far from significant (all *ps* > .2) and were therefore not included in the models reported here. The resulting partial correlations between reinstatement and source accuracy, controlling for age, are reported in Table 3. In no case did reinstatement reliably predict source accuracy over and above the factor of age group (all *ps* > .09, max *r* = .25).

**Within-participant point-biserial correlation analysis.** In a companion analysis to the trial-wise linear mixed effects analyses, we computed point-biserial partial correlations between binary source memory outcomes (correct, incorrect) and hippocampal activity, item- and category-level reinstatement estimates, and neural differentiation separately for each participant. We first computed the partial correlation between source accuracy and each factor while controlling for each of the remaining factors. Fisher z-transformed correlation coefficients for each factor were then carried over to two-sample *t*-tests (two-tailed) to test whether the magnitude of the relationship with source accuracy reliably differed between young and older adults. For all trial bins, the magnitude of the correlations were unmodified by group (*ps* > .07). We repeated these analyses after combining the data from the two age groups to form a single vector. The resulting correlations were then submitted to two-sided one-sample *t*-tests against a zero null. As is illustrated in supplementary Fig 1, for scene trials, these analyses yielded relationships that were significantly greater than zero between source accuracy and category-level reinstatement (mean *r* = .29, *p* = 8.44 x 10^-6^) and hippocampal activity (mean *r* = .06, *p* = 7.98 x 10^-5^), as well as an unexpected negative relationship between source accuracy and scene differentiation at encoding (mean *r* = -.07, *p* = .015). The relationship between source accuracy and item-level reinstatement of scene information was not significant (mean *r* = .02, *p* = .194). Turning to face trials, source accuracy significantly covaried with retrieval-related hippocampal activity (mean *r* = .07, *p* = 6.82 x 10^-5^) and item-level reinstatement (mean *r* = .04, *p* = .016). The mean correlations between source accuracy and category-level face reinstatement (mean *r* = .08, *p* = .056) and neural differentiation (mean *r* = .02, *p* = .663) were not significant.

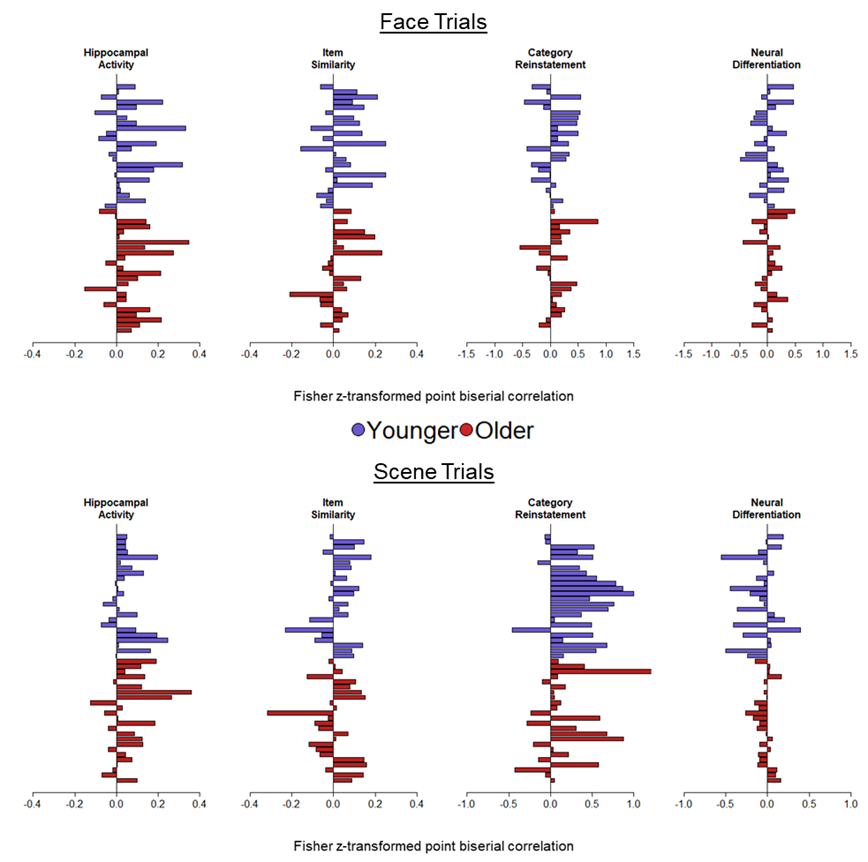

**Supplementary Figure 1 Point biserial correlation coefficients**

**Supplementary Table 1.** Participant demographics and *M (SD)* scores from the neuropsychological test battery

|  | Younger Adults | Older Adults | p-value |
| --- | --- | --- | --- |
| Years of Education | 15.46 (2.65) | 16.71 (2.44) | NS |
| MMSE | 29.25 (0.90) | 29.33 (0.70) | NS |
| CVLT Short Delay – Free | 13.75 (2.00) | 11.88 (2.86) | <0.01 |
| CVLT Short Delay – Cued | 13.83 (2.32) | 13.08 (2.15) | NS |
| CVLT Long Delay – Free | 14.13 (2.11) | 12.79 (2.62) | NS |
| CVLT Long Delay – Cued | 14.38 (1.93) | 13.46 (2.13) | NS |
| CVLT recognition – Hits | 15.71 (0.46) | 15.25 (1.07) | NS |
| CVLT recognition – False alarms | 0.33 (0.70) | 1.67 (1.61) | <0.01 |
| Logical Memory I | 33.00 (4.76) | 28.00 (4.11) | <0.01 |
| Logical memory II | 32.00 (4.80) | 25.83 (5.49) | <0.01 |
| SDMT | 62.33 (11.27) | 49.29 (7.91) | <0.01 |
| Trails A (s) | 20.20 (5.26) | 25.11 (6.46) | <0.01 |
| Trails B (s) | 44.12 (10.18) | 62.48 (16.77) | <0.01 |
| Digit Span Total | 19.71 (4.14) | 18.79 (3.49) | NS |
| Category fluency | 23.71 (4.91) | 22.46 (5.35) | NS |
| F-A-S | 49.17 (12.85) | 46.29 (12.75) | NS |
| WTAR | 42.42 (3.46) | 44.54 (4.06) | NS |
| Raven’s | 11.04 (0.86) | 9.50 (1.89) | <0.01 |
